# Comprehensive multi-omics characterization of gut microbiome extracellular vesicles reveals a connection to gut-brain axis signaling

**DOI:** 10.1101/2022.10.28.514259

**Authors:** Salma Sultan, Basit Yousuf, JuDong Yeo, Tamer Ahmed, Nour Elhouda Bouhlel, Heba Hassan, Zoran Minic, Walid Mottawea, Riadh Hammami

## Abstract

Microbiota-gut-brain axis is an evident pathway of host-microbiota crosstalk that is linked to multiple brain disorders. Microbiota released extracellular vesicles (MEVs) has emerged as a key player in intercellular signaling in host microbiome communications. However, their role in gutbrain axis signaling is poorly investigated. Here, we performed a deep multi-omics profiling of MEVs content generated ex vivo and from stool samples in order to get some insights on their role in gut-brain-axis signaling. Metabolomics profiling identified a wide array of metabolites embedded in MEVs, including lipids, carbohydrates, amino acids, vitamins, and organic acids. Interestingly, many neurotransmitter-related compounds were detected inside MEVs, including arachidonyl-dopamine (NADA), gabapentin, glutamate and N-acylethanolamines. Next, we aimed to identify commensal microbes with psychobiotic activity. We isolated 58 *Bacteroides* strains assigned to four genera, 11 species, and 4 new species based on 16S rDNA sequencing. We performed whole genome sequencing of 18 representative isolates, followed by a comparative analysis of the structure of polysaccharide utilization loci (PUL) and glutamate decarboxylase (GAD), a genetic system involved in GABA production. Quantifying GABA was done using competitive ELISA, wherein three isolates (*B. finegoldii, B. faecis*, and *B. caccae*) showed high GABA production (4.5-7 mM range) in supernatant whereas 2.2 to 4 uM GABA concentration was detected inside microvesicles extracted using ultracentrifugation. To test the biodistribution of MEVs from the gut to other parts of the body, CACO-2, RIN-14 B, and hCMEC/D3 cells showed a capacity to internalize labeled MEVs through an endocytic mechanism. Additionally, MEVs exhibited a dose dependent paracellular transport through CACO-2 intestinal cells and hCMEC/D3 brain endothelial cells. In vivo results showed biodistribution of MEVs to liver, stomach and spleen. Overall, our results reveal the capabilities of MEVs to cross the intestinal and blood brain barriers to deliver their cargoes of neuroactive molecules to the brain as a new signaling mechanism in microbiota-gut-brain axis communications.

## 1. Introduction

Appreciable evidence suggests the connection between microbiota-gut-brain-axis and the host’s brain activity ^1,2^. For instance, germ free mice exhibited a hyperactive hypothalamus-pituitary axis and a higher levels of stress hormones compared to conventional mice^3^. Also, gut microbiota have been shown to regulate emotional behaviors, modulates anxiety and memory processing^4–6^. Some probiotics known as “psychobiotics” have psychotropic like activities and are suggested to modulate mental and behavioural disorders. While some studies reported mitigated efficacy^7,8^, several trials support a role for psychobiotics in the normalization of brain processes related to stress responses and in mood improvements^9–12^. However, the mechanisms by which psychobiotics and gut microbiota interact with the gut–brain axis and modulate mental health remain hypothetical.

Microbiota released extracellular vesicles have been emerged recently as signaling molecules that mediate host-microbiota crosstalk^13^. MEVs are small membrane-bound phospholipid vesicles that range from 30 nm to 1 mm in size, with larger vesicles originating from the cell surface (microvesicles/ectosomes), and those on the smaller side being derived from either the plasma membrane or the endosomal system (exosomes)^14^. MEVs contain bacterial components such as nucleic acids, proteins, and cell wall components, and are involved in processes such as quorum sensing, biofilm formation, relief of environmental stresses, and host immunomodulation^15^. MEVs encase a spectrum of biologically active proteins, mRNA, miRNA, DNA, carbohydrates, and lipids, thus propagating the horizontal transfer of their cargo across both short and long distances^13^. The production and role of MEVs released by probiotic and commensal microbes in the gut environment are poorly investigated^16^, as MEVs have been predominantly examined in individual strains^15,17,18^. Previously, Kang et al.^19^ reported an important shift in stool MEV composition compared to the microbiome in an IBD DSS mice model. Moreover, same authors observed an attenuating effect of *Akkermansia muciniphila*-derived MEVs on the colitis severity^19^, suggesting the potential of MEVs as a biomarker and therapeutic agent. The recent report on increased levels of systemic LPS-positive bacterial extracellular vesicles in patients with intestinal barrier dysfunction provides some evidence on the capacity of MEVs to reach the systemic circulation^20^, and deliver and elicit a variety of immunological and metabolic responses in different organs including the brain. Therefore, MEVs should be considered of utmost importance delivery vehicles for host modulating metabolites, and thus can be regarded as promising psychobiotic candidates compared to clinical and regulatory limitations faced by fecal transplantation^21,22^. Harnessing the MEVs production ability of gut microbiota could decipher interaction with the gut–brain axis. In the current study, we postulate that microbiota interplay with the gut–brain axis involves microbiota extracellular vesicles (MEVs) as a communication-based mechanism. We investigated MEVs generated ex vivo and from stool samples of healthy adults as a potential cargo mechanism by which gut microbiota may exert psychobiotic effects.

## 2. Materials and methods

### 2.1. Isolation of MEVs from stool and ex vivo developed microbiota

The microbiota community from healthy donors (n=6 males and 6 females) were developed in an ex vivo model mimicking the large intestine^23^. The donors have not received antibiotics or probiotic supplementation for at least 3 months before the donation. Immobilization of microbiota was done by inoculating fecal samples into gellan (2.5 %, w/v) and xanthan (0.25 %, w/v) gum beads under anaerobic conditions as described previously^23^. Gel beads (30%) were transferred into a stirred glass reactor containing fresh Macfarlane culture medium. The fermentation conditions have been described before^23^. A total of 100 ml of the bioreactor culture after 15 days post-inoculation (microbiota stabilization period) was centrifuged at 10,000g for 30 min at 4°C. The supernatant was sterilized by filtration through 0.22 μ filter. MEVs were then isolated from the sterile supernatant by ultracentrifugation at 100,000g for 70 min. MEVs size was determined using a Malvern Zetasizer Nano ZS. The morphology of the isolated MEVs was inspected using transmission electron microscopy (TEM) as described previously^24^. Simply, the isolated MEVs in PBS was diluted 1:100 and fixed in 2.5% glutaraldehyde in a 0.1LJM sodium cacodylate buffer. The pellets were vortexed thoroughly and 10 μl of the suspension was added on to a carbon grid. Five minutes later, the grid was washed by a drop of distilled water. After drying, 10 μl uranyl less stain was added to the grid and incubated for 1 min before removing the excess stain and drying at room temperature before imaging with TEM.

### 2.2. DNA extraction for microbiome characterization through16S rRNA sequencing

The DNA was extracted from stool or 2 ml sample of the 15 days old ex vivo microbiome by FastDNA Spin Kit (MO BIO Laboratories Inc.) as per manufacture instructions with adding extra cycle of mechanical homogenization and 5 minutes of cooling on ice in between the 2 cycles. The extracted DNA was quantified using a Qubit fluorometer (Invitrogen; Carlsbad, CA, USA), it’s quality was checked by bioanalyzer 2100 (Agilent), and stored at −20 °C until used for further analysis.

### 2.3. DNA Library preparation for 16S rRNA sequencing

The microbiome diversity were determined by sequencing the V3-V4 regions of the 16S rRNA gene using the Illumina MiSeq platform (N*u*GUT Laboratory), using Illumina standard protocol as described in^23^. The raw paired end sequences fed to the Quantitative Insights Into Microbial Ecology pipeline 2 (QIIME 2)^25^ for quality preprocessing and determination of microbial composition and diversity indices.

### 2.4. Extraction of metabolites from MEVs

To minimize the contamination of soluble metabolites, 500 μL MEVs diluted in PBS were resuspended in PBS (7 mL) and re-centrifuged at 100,000*g* for 70 min (4°C) to wash it from the traces of media. The tube containing MEVs at the bottom was kept upside down for a few minutes to remove the remaining PBS. After that, 990 μL of LC/MS grade methanol and 10 μL of ^13^C-phenylalanine (1 mg/mL, internal standard) were added to the tube containing dried MEVs and they were gently mixed using a pipette to extract metabolites in MEVs. The result was transferred to the 1.5 mL Eppendorf tube and the insolubilized particles such as proteins formed in the extraction process were separated through centrifugation for 5 min at 16,000*g*. Then, the clear supernatant was passed through a 0.2 um polytetrafluoroethylene (PTFE) syringe filter (Millipore Corporation, Billerica, MA) to remove the remaining fine particle, followed by the evaporation of the solvent under vacuum using SpeedVac concentrator, and the dried samples were stored at -20°C until used for mass spectrometry analysis.

### 2.5. nLC-nESI-MS/MS analysis

The extracted metabolites from gut microbiota-derived MEVs were analyzed by using a nano liquid chromatography coupled on-line with nanoelectrospray ionization and mass spectrometry (nLC-nESI-MS/MS) (Thermo Fisher Scientific, Waltham, Massachusetts, US). The metabolites were separated on an EASY-Spray™ HPLC Columns (Thermo Scientific) with 15 cm × 75 μm ID, C18, 3 μm, 100 Å by using a water/acetonitrile/0.1% formic acid gradient. Samples (1μL) were injected by the autosampler and they were loaded onto the column for 60 min at a flow rate of 0.23 μL/min. The metabolites were then separated using a linear gradient from 10 to 100% of acetonitrile for 35 min, and kept for 10 min, followed by a gradient from 100 to 0% of acetonitrile for 5 min and again increased into 10% of acetonitrile for 10 min.

Eluted metabolites were directly sprayed and transferred into a mass spectrometer by electrospray ionization (ESI). Full-scan MS spectra (m/z 100–1000) were acquired at a resolution of 70,000. The automatic gain control settings were 3 × 10^6^ for full MS and 1 × 10^5^ for MS/MS scans and the fragmentation of molecules occurred in the collision-induced dissociation (CID) cell in the linear ion trap. Precursor ions were separated using a 2 m/z isolation window and fragmented with a normalized collision energy of 35%.

### 2.6. *nLC-nESI-MS/MS* Data processing and analysis

Mass spectrometry data were processed and analyzed by MS dial version 4.60 along with Massbank and Human metabolome database (HMDB) to facilitate the identification of target molecules. Default parameters were used for the identification of individual compounds in samples, and contaminations detected in the blank were excluded from the list of identified compounds.

### 2.7. Characterization of commensals with psychobiotic activity

#### 2.7.1. Bacterial strains isolation

Fecal samples were collected from an adult healthy Canadian women donor after taking consent from the University of Ottawa Research Ethics Board and Integrity protocol and the donor. The fecal sample was homogenized with reduced peptone water (20%, w/v). Serial dilution of the samples was done in PBS (10^−1^ to 10^−8^) and then spread on Fastidious anaerobic agar with 0.5% yeast extract (FAAy) and Brain heart infusion with yeast extract, cysteine, and hemin (BHIych) and allowed to grow under anaerobic conditions (5% hydrogen, 10% CO2, 85% nitrogen) for five days. Colonies were picked up and streaked separately on respective agar plates. Single colonies from each plate were inoculated into culture broth for preservation in 25% glycerol at - 80 °C.

#### 2.7.2. Glutamate Decarboxylase assay

Preliminary screening to select *Bacteroides* isolates for molecular identification was performed based on GAD assay ^26^. We prepared a hypertonic test substrate solution consisting of 0.1 g of L-glutamic acid, 30 ul Triton X-100, 9 g NaCl, 0.005 g Bromocresol green as an indicator in 100 ml of sterilized water with pH of 4.0, which gives the solution yellow coloration. Bacterial culture (2 ml) grown for 48 h was centrifuged, the pellet was washed with 0.9% saline, and 0.3 mL test solution was added to the washed pellet. Color changes from yellow to green/blue were considered positive for GAD activity.

#### 2.7.3. DNA extraction, 16S rDNA-based characterization, and taxonomic assignment

DNA was extracted from isolates using a NucleoSpin Microbial DNA kit (Macherey-Nagel, Duren, Germany) following the manufacturer’s instructions. PCR was performed to amplify the 16S rRNA gene using universal primers 8F and 1391R ^26^. PCR reaction mixture consisted of 1× PCR buffer, 1.5 mM MgCl2, 0.2 mM NTPS (Invitrogen, Carlsbad, CA, USA), 1 μM of each primer, 1U of Taq DNA polymerase (Invitrogen), and 50-100 ng of bacterial DNA in 50 μl of nuclease-free water. PCR amplification program included one cycle of 95 °C for 3 min; 30 cycles of 95 °C for 30 s, 55 °C for 30 s, 72 °C for 60 s; and finally, one cycle of 72 °C for 5 min. PCR amplification was confirmed using gel electrophoresis (1.2% agarose gel with ethidium bromide). QIAquick PCR purification kit (Qiagen, Hilden, Germany) was used to clean up the PCR products, followed by sequencing at Ottawa Hospital Research Institute’s DNA sequencing facility (Ottawa, ON, Canada). Quality assessment of the Chromatograms of the sequences was analyzed using BioEdit program. The taxonomic attribution was done with RDP classifier ^27^, and a nucleotide similarity cut-off of 99% (16S rRNA gene) was used to identify an isolate to the species level ^28^.

#### 2.7.4. Phylogenetic Analysis

For 16S rRNA, *gadA*, and *gadB* genes, phylogenetic trees were constructed using the Molecular Evolutionary Genetics Analysis (MEGA11) ^29^. Initial tree(s) for the heuristic search were obtained automatically by applying Neighbor-Join and BioNJ algorithms to a matrix of pairwise distances estimated using the Jukes-Cantor model and then selecting the topology with superior log likelihood value. Bootstrap resampling was performed with 500 bootstrap replications to have the confidence of tree topologies. The percentage of trees in which the associated taxa clustered together is shown next to the branches.

#### 2.7.5. Quantification of SCFAs Using Gas Chromatography

Production of SCFAs such as acetate, butyrate, and propionate were quantified using gas chromatography coupled with the flame ionization detector (GC-FID) (Shimadzu GC-2030) ^26^. For sample preparation, 2 mL bacterial cultures were centrifuged twice at 14,000 g for 30 min at 4 °C and sterilized with a 0.22-μm syringe filter. 2-ethyl butyric acid was added to each sample at a concentration of 0.5 mM as an internal standard. Data analysis was done using Lab Solutions software developed by Shimadzu Corporation, Japan. Samples were analyzed in duplicates, and results were expressed in mM.

#### 2.7.6. Whole Genome Sequencing, annotation, and comparative genome analysis

Library for whole genome sequencing was prepared using Nextera™ DNA Flex Library Prep (Illumina; San Diego, CA, USA) as per the recommended protocol. Paired end sequencing (2 × 151 bp) was performed using MiSeq Reagent Kit v2 with Illumina MiSeq platform (N*u*GUT Research Platform, University of Ottawa, Ottawa, ON, Canada). De-novo assembly of the generated reads was done using Velvet Assembler V1.0.0 incorporated in Illumina BaseSpace Sequence Hub (Illumina). The assembled contigs were annotated using Rapid Annotation using Subsystem Technology (RAST) server ^30,31^. GAD system genes were binned together to determine the structure of the GAD system in different *Bacteroides* species. The number and structure of PULs in each genome were determined by automatic prediction using PULpy software ^32^.

#### 2.7.7. Microvesicle purification and metabolite extraction

One hundred twenty milliliters of a 72h culture of 4 *Bacteroides* strains (*B. finegoldii, B. faecis, B. caccae*, and *Bacteroides*) was centrifuged at 15,000 g at 4°C. The supernatant was filtered 0.22 μm filter; consequently, ultracentrifugation was performed at 100,000 g for 1 h (Optima L-90K ultracentrifuge; Beckman Coulter). The supernatant was stored at -20°C, and the pellet was washed with 5 mL of sterile PBS, followed by one more ultracentrifugation. The vesicle pellet was resuspended in distilled water.

For metabolite extraction, 900 μl of methanol (−20°C) was added to 100 μl of the resuspended OMVs and then kept at −80°C for 15 min. The sample was thawed for 3 min at room temperature and then thoroughly vortexed again, followed by centrifugation for 30 min at 16,000 g at 4°C. Vacuum drying of the supernatant was performed using a SpeedVac concentrator (Thermo Fisher Scientific™, USA). The dry extract was then dissolved in 100 μl of sterile water and then was analyzed for GABA quantification along with supernatant using ELISA.

#### 2.7.8. ELISA tests for quantification of GABA

Single colonies were inoculated into FAB and allowed to grow for 24 hours. The preculture was then added 1.25% (v/v) to 8 mL FAA supplemented with 10 mM glutamate (pH = 6.5) and incubated at 37°C for 72 h under anaerobic conditions. 2 ml of bacterial cultures were centrifuged at 14,000 × g, 10 min, 4°C and supernatants were collected and filtered through 0.22 μm filter and were aliquoted and stored at −20 °C until used for ELISA assays. GABA was quantified in three independent replicates using an ELISA kit (Abnova, Taipei, Taiwan), according to the manufacturer’s protocols.

#### 2.7.9. Statistical Analysis

The statistical analysis was carried out by GraphPad Prism 8.0 (GraphPad Software Inc., CA, USA).

### 2.8. Cell Culture Experiments

Unless specified, all reagents were purchased from MilliporeSigma (Toronto, ON). Dulbecco’s modified eagle medium (DMEM), RPMI-1640 media, L-glutamine, penicillin/streptomycin, chemically defined lipid concentrate, HEPES, trypsin-ethylenediaminetetraacetic acid (trypsin-EDTA), trypan blue, Dulbecco’s phosphate-buffered saline (DPBS) were purchased from ThermoFisher Scientific (Gibco, Nepean, ON). Tissue culture well plates (12, 24, and 48), flasks (25 and 150 cm2), and inserts (0.4μm, 12 well plates) were purchased from Corning (Maine, USA). Tissue Culture Inserts, 0.4 μm (12 well plates) were purchased from (ThinCert, Greiner Bio-One, Monroe, NC). Rat collagen I and recombinant human basic fibroblast growth factor (FGF-2) were purchased from R&D Systems (Toronto, ON). Endothelial cell growth basal medium-2 (EBM-2) was purchased from Lonza (Kingston, ON).

#### 2.8.1. Endocytosis activity

RIN-14 cells (ATCC® CRL 2059) were grown to confluence in complete culture media (CCM) constituted of RPMI-1640 supplemented with 10% heat-inactivated FBS (HI-FBS), 100U/mL penicillin, 100 mg/mL streptomycin^33^. CACO-2 cells (ATCC HTB 37) were grown to confluence in CCM constituted of DMEM supplemented with 10% HI-FBS, 100U/mL penicillin, 100 mg/mL streptomycin^34^, while hCMEC/D3 (Cedarlane labs, CLU 512) were grown on collagen-coated flask using CCM constituted of endothelial basal medium (EBM-2) supplemented with 5% HI-FBS, 100U/mL penicillin, 100 mg/mL streptomycin, hydrocortisone (1.4μM), ascorbic acid (5μg/mL), chemically defined lipid concentrate (1%), HEPES (10mM), basic fibroblast growth factor (1 ng/mL)^35^. All cells were maintained at 37 °C in a 5% CO2 humidified incubator (VWR, Mississauga, Ontario). For subculturing, different cells were dissociated using 0.05% trypsin–EDTA, centrifuged, and washed 2 times using DPBS (pH 7.4) to remove traces of trypsin^36^.

CACO-2 at a density of 2×105 cells^37^ and RIN-14B at a density of 2.5×105 cells were seeded on 12 well plates^38^. In contrast, hCMEC/D3 cells were seeded at a density of 2×105 on a collagen-coated 12 well plate using rat tail collagen (150 μg/mL)^39^ and maintained at 37 °C in a 5% CO2 humidified incubator for 2 days (CACO-2 and hCMEC/D3 cells) and 5 days (RIN-14B) with refreshing media every 3-4 days. Microbiota extracellular vesicles (MEVs) equivalent to 2 grams of stool were labeled using Cyanine 7 (Cy7), added to each well of different cells, and incubated for 24 hrs. Conditioning media containing the unbound MEVs was discarded. Cells grown in the absence of labeled MEVs served as a negative control. The cells were washed three times, mounted in DPBS, and then imaged using an inverted Zeiss AxioObserver 7 microscope (Carl Zeiss Canada Ltd, Toronto, ON).

### 2.8.2. Transport activity

#### 2.8.2.1. CACO-2 cells

CACO-2 cells were seeded at a density of 5×104 (P8) onto ThinCert™ tissue culture inserts (12 well plates, 0.4 μm, 108 pores, 1.131 cm2) and then maintained in CCM^40^. Culture media were refreshed every 2 days. Cells were allowed to grow and differentiate for 28 days. On day 28, CCM was replaced by phosphate-buffered saline (PBS), and resistance values (Ω) were measured for all tissue culture inserts, including blank, using Millicell ERS-II voltohmmeter^41^ (MilliporeSigma, Toronto, ON). The TEER values for each insert were calculated according to the following equation:

TEER = (R (Cells) - R (Blank)) x tissue culture inserts surface area (Ω.cm2) ^41^ MEVs labeled with fluorescein isothiocyanate (FITC)^42,43^ (equivalent to 0.9, 1.8-, and 3.6-grams stool) were suspended in CCM and added to the apical side of the tissue culture insert. Fresh CCM was added to the basolateral side, then conditioned culture supernatant was collected from the basolateral side at 0, 3, and 24 hrs. The fluorescence intensity was measured using a TECAN plate reader at an excitation/emission wavelength of 485/535 nm. PBS replaced conditioned media in the apical and basolateral compartments, and TEER values were reported for various treatment groups.

#### 2.8.2.2. hCMEC/D3 cells

For the blood brain barrier transport study, hCEC/D3 cells were seeded at a density of 5×104 onto collagen-coated inserts (12 well plates)^35^ using CCM and allowed to differentiate for 4 days. TEER resistance values were recorded every day. The TEER values for each insert were calculated according to the aforementioned equation. After identifying the time point showing the peak TEER value, MEVs labeled with Cy7^44^ (equivalent to 0.9, 1.8-, and 3.6-grams stool) were suspended in CCM and added to the apical side of the tissue culture insert (Corning). Fresh CCM was added to the basolateral side, then conditioned culture supernatant was collected from the basolateral side at 0, 3, and 24 hrs. The fluorescence intensity was measured using a TECAN plate reader at an excitation/emission wavelength of 743/780 nm. PBS then replaced conditioned media in the apical and basolateral compartments, and TEER values were measured for various treatment groups.

### 2.8.3. Cell Culture Statistical Analysis

To compare the mean values of different results, one-way ANOVA (SPSS, version 23) followed by LSD as a post hoc test was used to determine the statistically significant difference for data involving more than 2 experimental groups. Paired sample t-test was used to identify the statistically significant difference between CACO-2 and hCMEC/D3 cells’mean TEER values before and after MEVs treatment compared to the control.

Independent sample t-test was used to compare the data of the in vivo experiments. All results were expressed as Mean ± Standard Error. All the figures were designed using Microsoft 365® Apps for enterprise (Excel and Power Point) and Adobe Photoshop CS (version 8).

### 2.9. In vivo assessment of MEVs biodistribution

To assess the in vivo trafficking of MEVs to the systemic circulation and their potential diffusion across the blood-brain barrier, the inbred strain C57BL/6 mice were used as an animal model. All *in vivo* experiments were conducted following the approved animal protocol (HSe-3857) by the University of Ottawa Animal Care Committee and according to the guidelines of the Canadian Council on Animal Care (CCAC). All animal protocols comply with the NIH Guide for Care and Use of Laboratory Animals (Animal Welfare Assurance # A5043-01). A total of 9 C57BL/6 mice (Female, 8-12 weeks, 16-21 grams) were used for the different experimental groups. Mice were acclimatized for 3-7 days under ambient conditions (12 hours light/ dark cycle at 22 ±2 °C and humidity 50-60%) with free access to sterile water and a standard rodent soft chow ad libitum. The experimental group included 6 mice (female, n=6), were injected through the tail vein with purified MEVs labeled with Cy7 (equivalent to 12 g stool), while the control group included 3 mice (n=3) and injected with an equal volume of phosphate buffer saline (PBS; placebo control). At 24 hours, trunk (heart) blood was collected from intubated animals (1-3 isoflurane) to analyze serum fluorescence levels. All mice were then euthanized to collect different tissues, including liver, fat, muscle, spleen, large intestine, and brain, for organ imaging using the IVIS Lumina XR (Perkin Elmer, Woodbridge, ON) with excitation/emission wavelengths of 756/779 nm to assess the MEVs biodistribution. Fluorescence intensity of imaged organs collected from test and control mice was expressed as average radiant efficiency^45,46^.

### 2.10. GenBank submission and accession numbers

16S rRNA gene sequence data were deposited in the GenBank database with accession numbers OP690542-OP690599. Whole-genome sequence data is available in GenBank under the accession numbers ????????????.

## 3. Results

### 3.7. Physical characteristics of the isolated MEVs

Transmission electron microscopy (TEM) and Zetasizer showed that the isolated MEVs are intact, and membrane enclosed and their size falls in the range of exosomes, microvesicles and bacterial outer membrane vesicles^47^ (Figure 1A-B).

**Figure 1:**
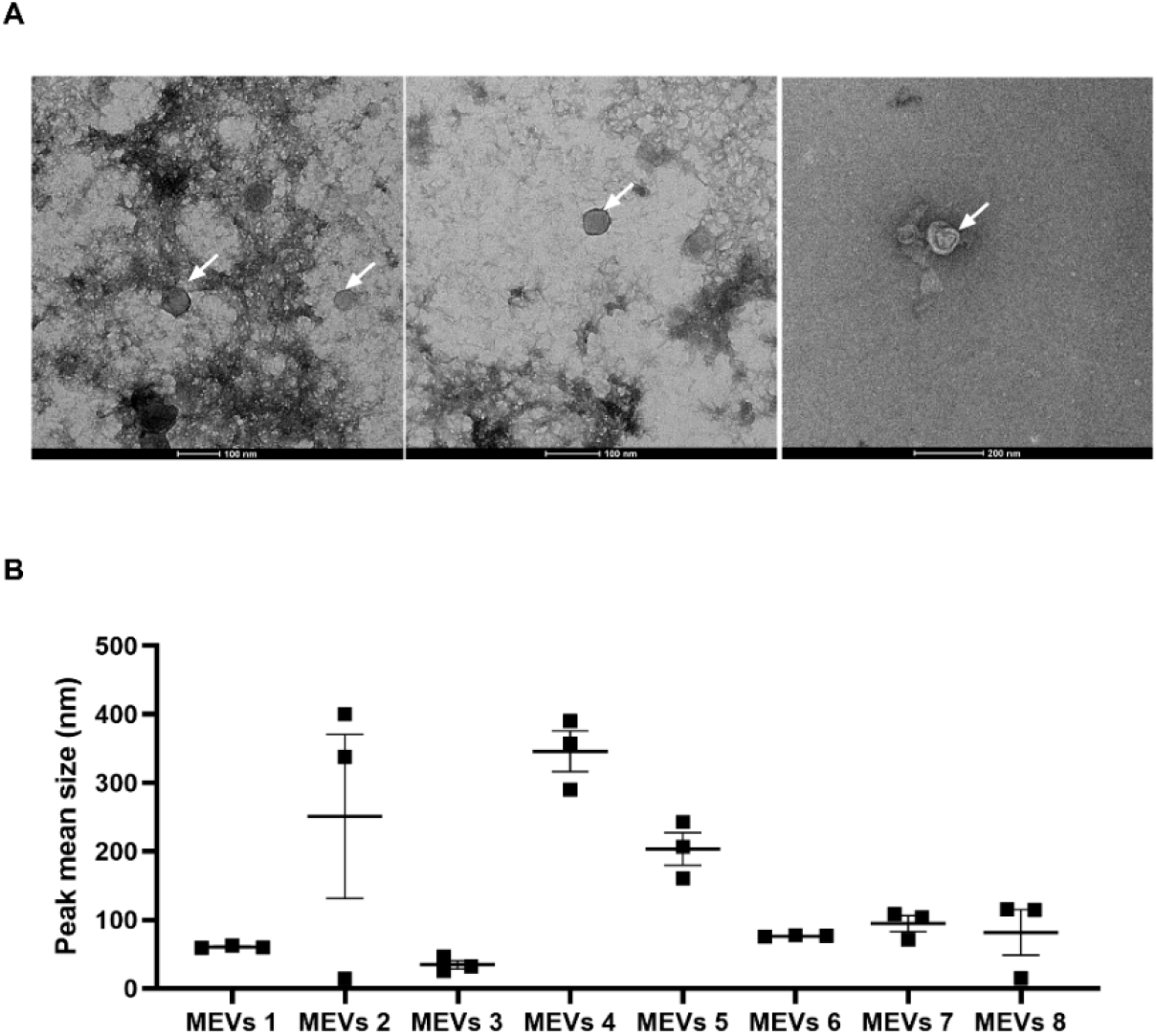
Transmission electron microscopy (TEM) and size of isolated vesicles. TEM of the isolated vesicles. **(B)**Average Size ±SEM of the isolated MEVs from 8 samples as determined by a Malvern Zetasizer Nano ZS. Horizontal lines represent Mean ±SEM.

### 3.8. Metabolic content of MEVs determined by nLC-ESI-MS/MS analysis in the <u>positive mode</u>

The identified lipid species in gut microbiota-derived MEVs in the positive mode are summarized in **Figure 2**. More than 40 lipid species were identified in the isolated MEVs from the stool and fermented samples in the positive mode. The major lipid classes found in the samples were spingoid bases (SPB), acyl carnitines (CAR), lyso-phosphatidylethanolamine (LPE), and lyso-phosphatidylcholine (LPC), possessing only one acyl chain in their chemical structure. Given that a large portion of lipids species found in MEVs is the primary sources for the construction of lipid bilayer membrane of MEVs due to their amphiphilic nature, it is expected that the high proportion of lipids in the membrane of MEVs consists of single acyl chain lipids such as SPB, CAR, LPE and LPC (**Figure 2-b**). A few phospholipids such as phosphatidylcholine (PC) and phosphatidylethanolamine (PE) having two acyl chains were detected in the positive mode, but those are not included in **Figure 2** as it needs more data processing for identification. Overall, MEVs isolated from fermented samples showed a higher abundance in ion intensity compared to that of the stool sample.

**Figure 2.**
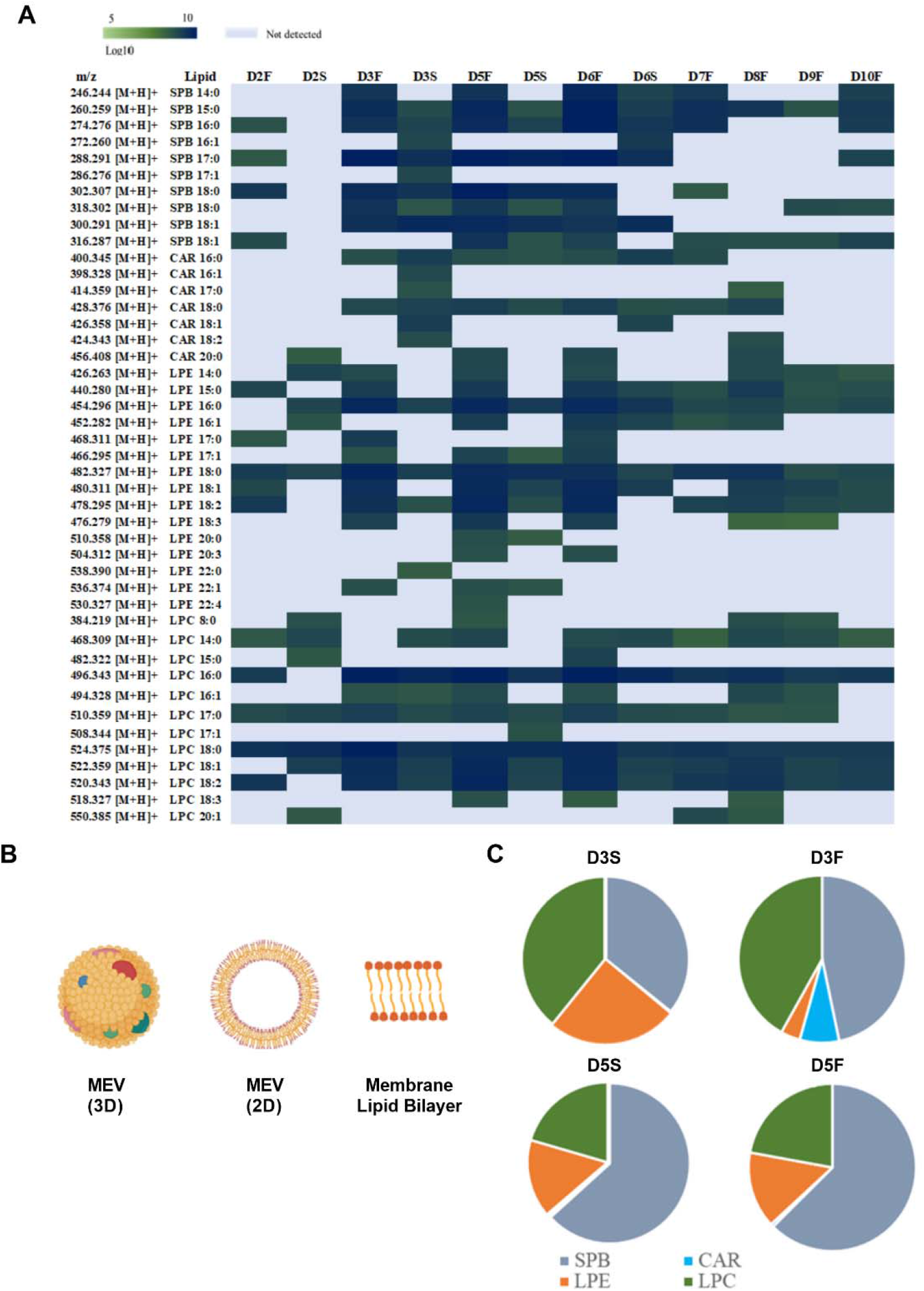
Mass spectrometry-based lipidomics in the determination of lipid profiles of gut microbiota-derived MEVs <u>in the positive mode</u>. The heat map (a) describes the relative abundances of individual lipid species found in isolated MEVs (D3F: Donor3-fermented; D3S: Donor3-stool, D5F: Donor5-fermented; D5S: Donor5-stool). The possible structure of MEV and its lipid bilayer membranes are provided in the (b) section. The pie charts (c) show the different ratios of lipid classes in the four samples. SPB; Sphingoid bases, CAR; Acyl Carnitines, LPE; Lyso-phosphatidylethanolamine and LPC; Lyso-phosphatidylcholine.

In **Figure 2-c**, the discrepancy in the ratio of lipid classes between stool an ex vivo samples is provided in a pie chart style. The MEVs from donor 3 and donor 5 showed a different ratio in lipid classes in both fermented- and stool samples. Donor 5 showed a higher proportion of SPB than that of LPE and LPC, displaying a different pattern with Donor 3 that has a less proportion of SPB. Interestingly, a similar ratio of lipid class was found between MEVs isolated from the same donor’s stool and fermented samples, especially donor 5, which may be due to similar microbiome composition within the two samples of the same donor, leading to the production of analogous MEVs.

The metabolite profiles, except lipid species, in MEVs isolated from stool and fermented samples were also determined in the positive mode and provided in **Figure 3**. A wide spectrum of molecules was identified including carbohydrates, amino acids and steroid derivatives in this limited data. **Figure 3** shows that among the metabolic content of MEVs is a set of neuroactive molecules including choline, gabapentin, phenylalanine, tyrosine, arachidonyl-dopamine (NADA), L-glutamic acid, and N-acylethanolamines.

**Figure 3.**
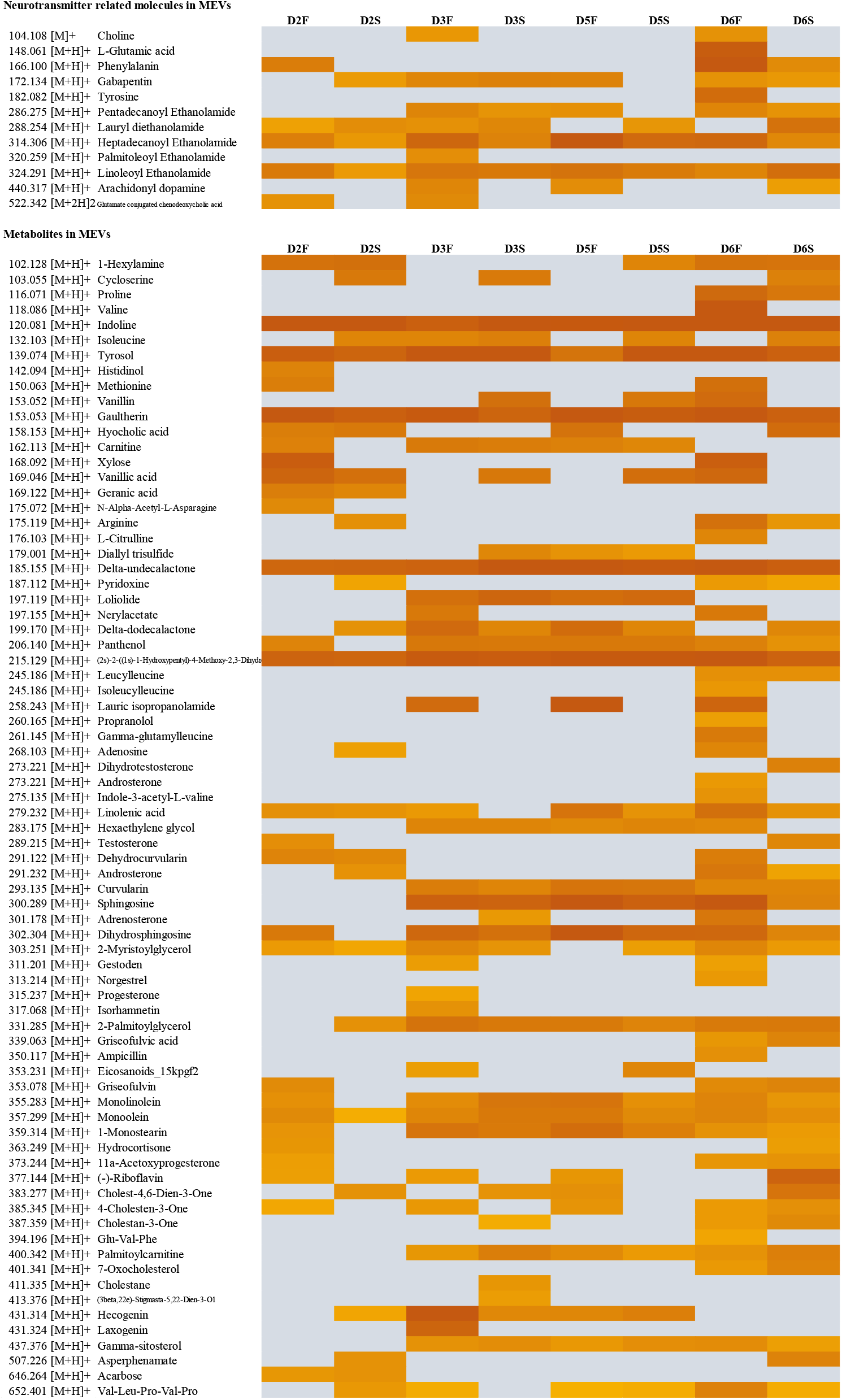
Mass spectrometry-based metabolomics in the determination of cargoes inside MEVs <u>in the positive mode</u>. The heat maps describe the relative abundances of individual metabolites identified in isolated MEVs.

### 3.9. Metabolic content of MEVs determined by nLC-ESI-MS/MS analysis in the <u>negative mode</u>

The lipid species identified in gut microbiota-derived MEVs in the negative mode are summarized in **Figure 4**. In total, 54 lipid species were identified in the isolated MEVs from the stool and fermented samples in the negative mode. The dominant lipid classes found in the MEVs were lyso-phosphatidylethanolamine (LPE), and several species of N-acetylglycine (NAGly), N-acyl glycylserine (NAGlySer), phosphatidylethanolamines (PE), and phosphatidylinositol (PI) were also identified in the negative mode. Overall, the different classes of lipid species were found in the negative mode compared to that of the positive mode, which might be due to the discrepancy in the ionization properties of each lipid class. Several lipid species that are conjugated with high unsaturated fatty acids (HUFAs) such as linoleic acid (18:3) and arachidonic acid (20:4) were identified in the negative mode.

**Figure 4.**
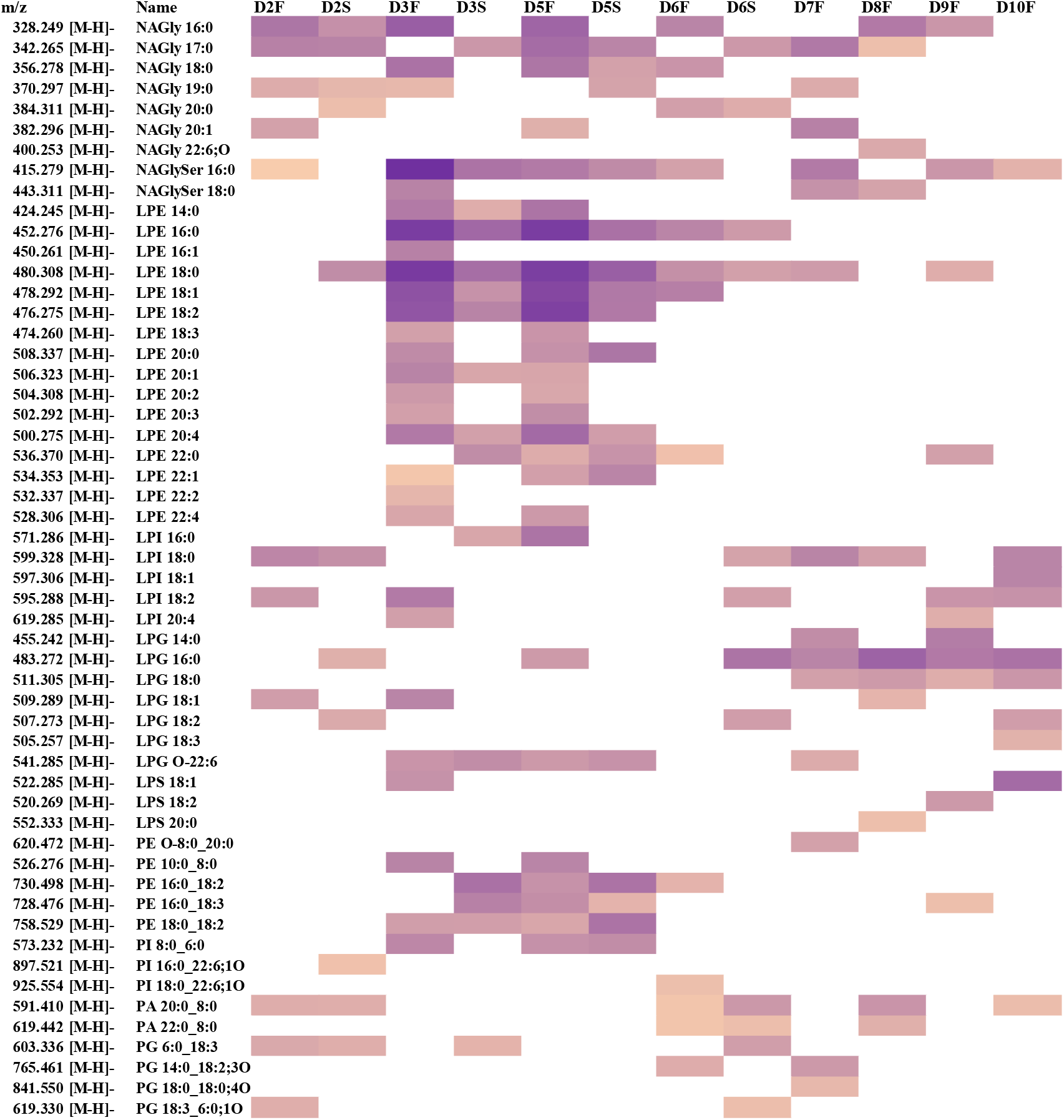
Mass spectrometry-based lipidomics in the determination of lipid profiles of gut microbiota-derived MEVs <u>in the negative mode</u>. The bar describes the relative abundances of individual lipid species found in isolated MEVs. NAGly: N-Acetylglycine; NAGlySer: N-acyl glycylserine; PE: Phosphatidylethanolamines; PI: Phosphatidylinositol.

The metabolite profiles, except lipid species, in MEVs isolated from stool and fermented samples were also determined in the negative mode and summarized in **Figure 5**. A wide range of metabolites was found in MEVs in the negative mode including carbohydrates, amino acids, vitamins, and organic acids, in particular, some neurotransmitter-related compounds were also identified. For instance, 3,4-dihydroxymandelic acid (DOMA), a metabolite of norepinephrine that is referred to as a representative neurotransmitter, was identified at m/z [M-H]^−^ 183.0 in the negative mode. Arachidonyl dopamine at m/z [M-H]^−^ 438.2, which is found in the positive mode, was also identified in the negative mode. Dopamine, a representative neurotransmitter in humans, was found in the gut microbiota-derived MEVs as a conjugated form by associating with arachidonic acid (20:4).

**Figure 5.**
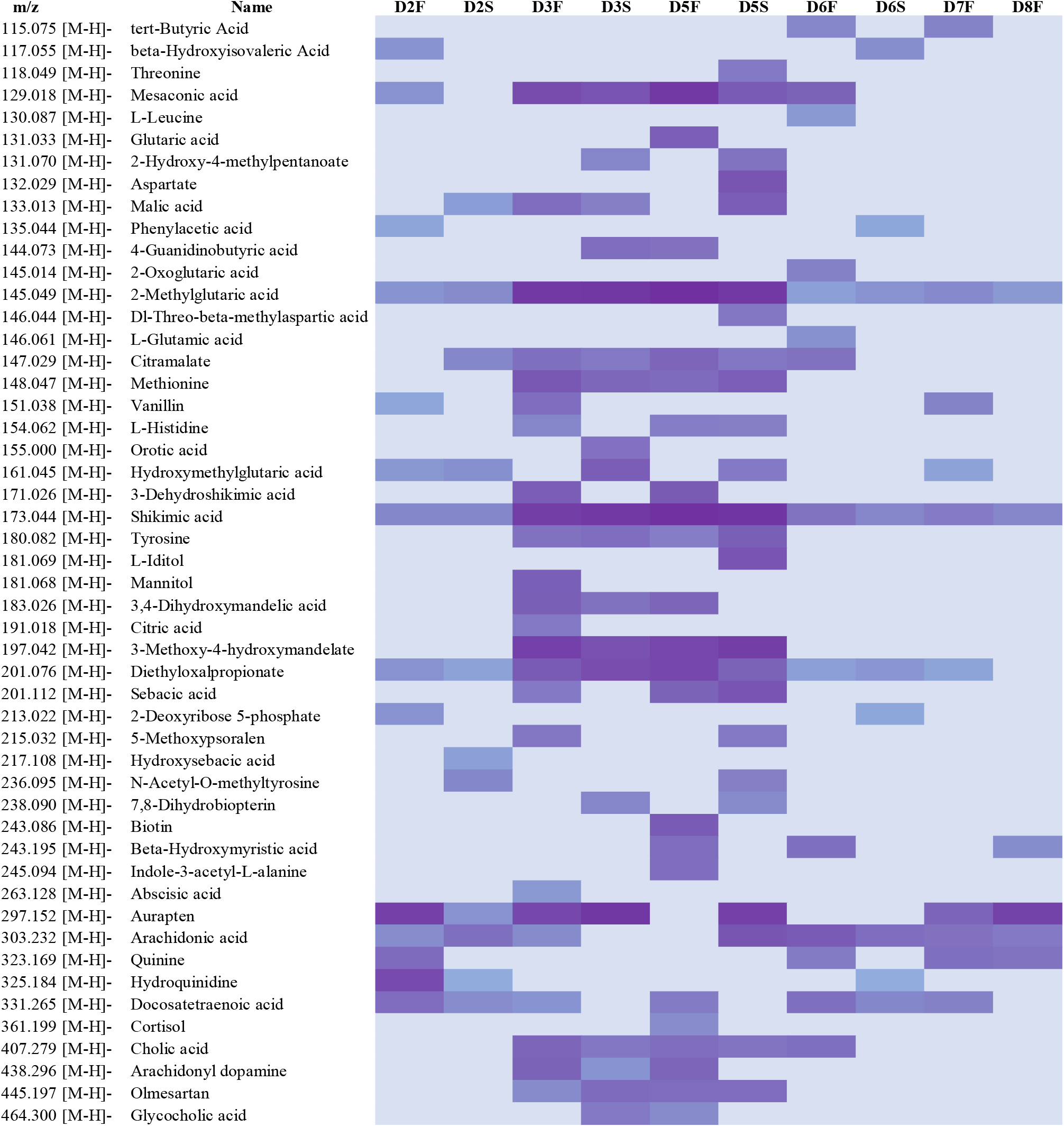
Mass spectrometry-based metabolomics in the determination of cargoes inside MEVs in the <u>negative mode</u>. The bar describes the relative abundances of individual metabolites identified in isolated MEVs.

### 3.10. Gut Microbiota Structure

We characterized the microbiota structure from which we extracted the MEVs via 16S rRNA sequencing. Stool samples showed higher diversity compared to the ex vivo developed microbiome (**Figure 6A**). The microbiota from stool and ex vivo samples followed the common structure of microbiome that is dominated with Bacteroidetes and Firmicutes with lower abundance of Proteobacteria and Actinobacteria and low percentages for other phyla (**Figure 6B**). The ex vivo-developed microbiome showed depletion of Ruminococcaceae and Lachnospiraceae along with expansion of Enterobacteriaceae and Veillonellaceae (**Figure 6B-C**) compared to the stool samples. still the effect of donor on the microbiome structure is evident where PCA analyses showed the samples clustered mainly by sample type (stool vs ex vivo) then by the donor (**Figure 6D**). 102 taxonomy features showed differential abundance between stool and *ex vivo* microbiota (Figure 6E, table 1). These features are dominated by *Faecalibacterium* spp. (046cb7522f5d3589a6dcbcd1f2cda95d), Ruminococcaceae family (70392e48fb80c7c9fd7adeae40a3585e) mainly *Gemmiger formicilis* (b1ffa316da920a0242cfa65f45395023) and Lachnospiraceae (0cfab271db3ec8f652b4159488aab51f) mainly the *Clostridium* genus (db70b8d89735474e9c237ba7be41c449).

**Table 1.** Number of isolates from each species.

| Bacteria | Number of Isolates |
| --- | --- |
| *Bacteroides cellulosilyticus* | 5 |
| *Bacteroides dorei* | 3 |
| *Bacteroides faecis* | 2 |
| *Bacteroides stercoris* | 9 |
| *Bacteroides uniformis* | 2 |
| *Bacteroides vulgatus* | 13 |
| *Bacteroides ovatus* | 1 |
| *B. Finegoldii* | 1 |
| *B. Zhangwenhongii* | 2 |
| *Parabacteroides johnsonii* | 3 |
| *Alistipes putredinis* | 1 |
| *Butyricimonas virosa* | 1 |
| *Bacteroides* spp. | 12 |

**Figure 6.**
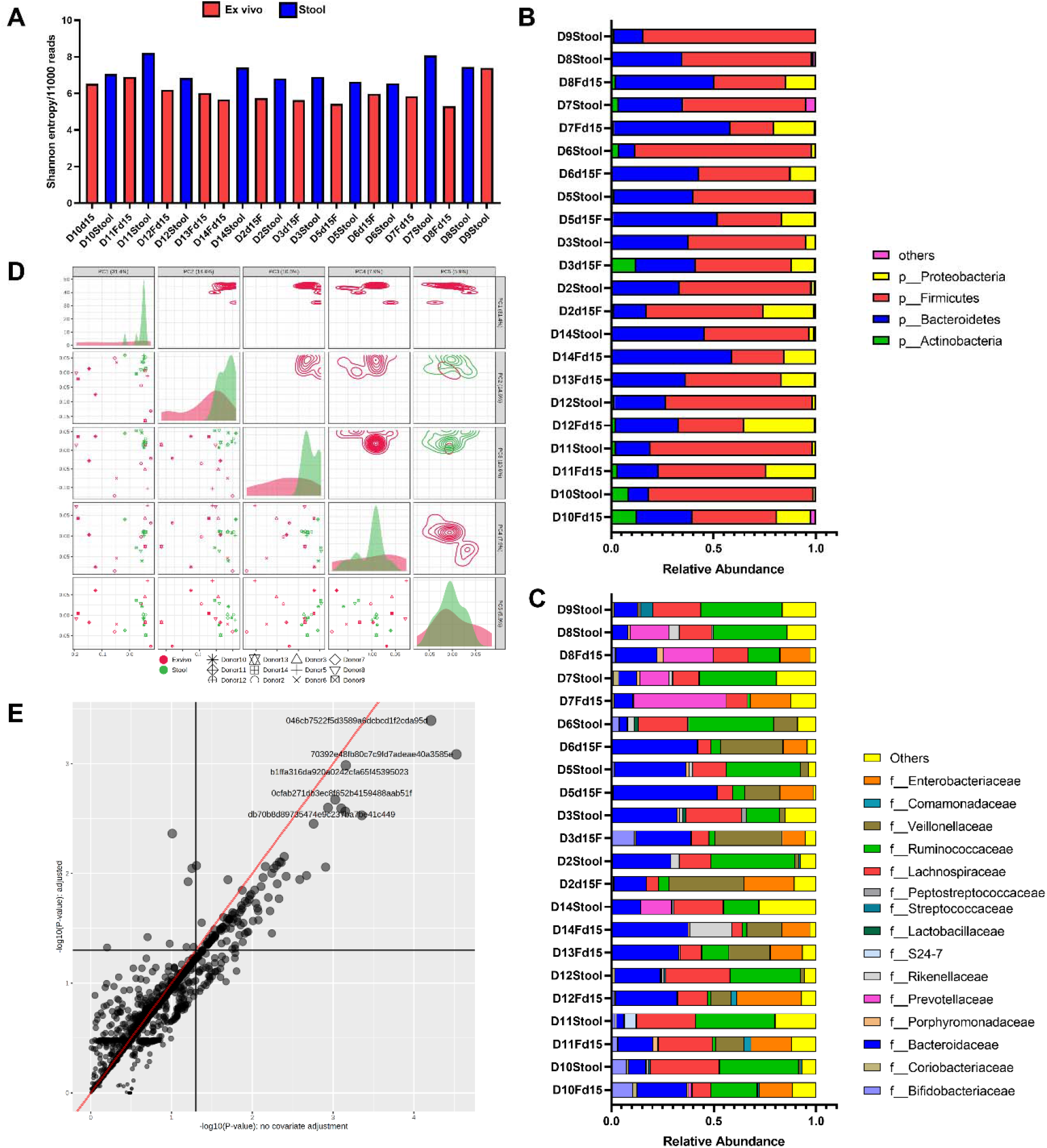
Gut microbiota structure from which MEVs were generated. (A) Diversity of gut microbiota in each sample as indicated by Shannon indices. (B) and (C) Composition of the gut microbiota at the phyla and family level. (D) Principal component analyses of different samples based on the relative abundance of different features identified in each sample generated by Metaboanalyst 5.0 using the features table generated by QIIME 2.6. (E) Linear model showing the variable features between stool and ex vivo generated microbiome using donor as a covariate for adjustment.

### 3.11. Isolation of Human Gut Bacteria and Selection and Identification of *Bacteroides* strains

For the isolation of gut bacteria, we picked up late appearing 130 colonies from the BHIych and FAAy plates after 3 to 5 days. Fifty-eight possible *Bacteroides* isolates were selected based on positive GAD assay and were followed for 16S rDNA based molecular identification. 16S rRNA gene sequences were analyzed for the nearest match in Ribosomal Database Project (RDP) which revealed 38 isolates belonged to *Bacteroides* genera, 5 isolates to *Parabacteroides* genera, 1 each to *Butyricimonas* and *Alistipes* (**Table 1**). Other 12 isolates showed highest nucleotide matching with uncultured bacteria and were then analyzed against type strains. Notably, we found only a single match, *B. massiliensis* in RDP for these 12 isolates with nucleotide similarity percentage varying from 98.2% to 99.2%. Other type strains showed less than 80% similarity indicating these 23 isolates to be possible new species (Edgar, 2018). Based on multiple sequence alignment, these 12 possible new species could be grouped into 3 *Bacteroides* species.

### 3.12. Characterization of GAD and PUL systems in *Bacteroides* strains

Whole genome sequencing was performed for 18 representative strains which included 10 *Bacteroides* species (*B. cellulosilyticus* UO.H1027; UO.H1030, *B. dorei* UO.H1033, *B. faecis* UO.H1051, *B. stercoris* UO.H1035; UO.H1039; UO.H2001, *B. uniformis* UO.H1043, *B. vulgatus* UO.H1015; UO.H1016, *B. ovatus* UO.H1053, *B. Finegoldii* UO.H1052, *B. Zhangwenhongii* UO.H1054, and *B. caccae* UO.H2003), 1 *Parabacteroides* species (*P. johnsonii* UO.H1047 and UO.H1049) and two new *Bacteroides* sp. UO.H1001; UO.H1004. In silico analysis of GAD genetic system was performed to see the difference in nucleotide similarity as well as structure amongst the 11 different species and how it correlated with difference in GABA production capabilities. In *B. Finegoldii, B. caccae* and *B. faecis, gadB* (glutamate decarboxylase) and *gadA* (glutaminase) genes were in close vicinity with only 11 or 9 bp distance between them (**Figure 7**). Notably, *gadC* (glutamate/GABA antiporter-encoding gene) and *gadD* (potassium (K+) channel transporter) were in close proximity almost in all *Bacteroides* strains, implying their functional importance. Notably, *B. finegoldii* UO.H1052 and *Bacteroides* sp. UO.H1001; UO.H1004 showed the absence of *gadC* and K+ channel genes possibly due to lack of annotation (**Figure 7**). Interestingly, *gadB* gene of *B. finegoldii* and *Bacteroides* sp. showed 88 and 94% nucleotide sequence similarity invoking their novel GAD genetic system.

**Figure 7.**
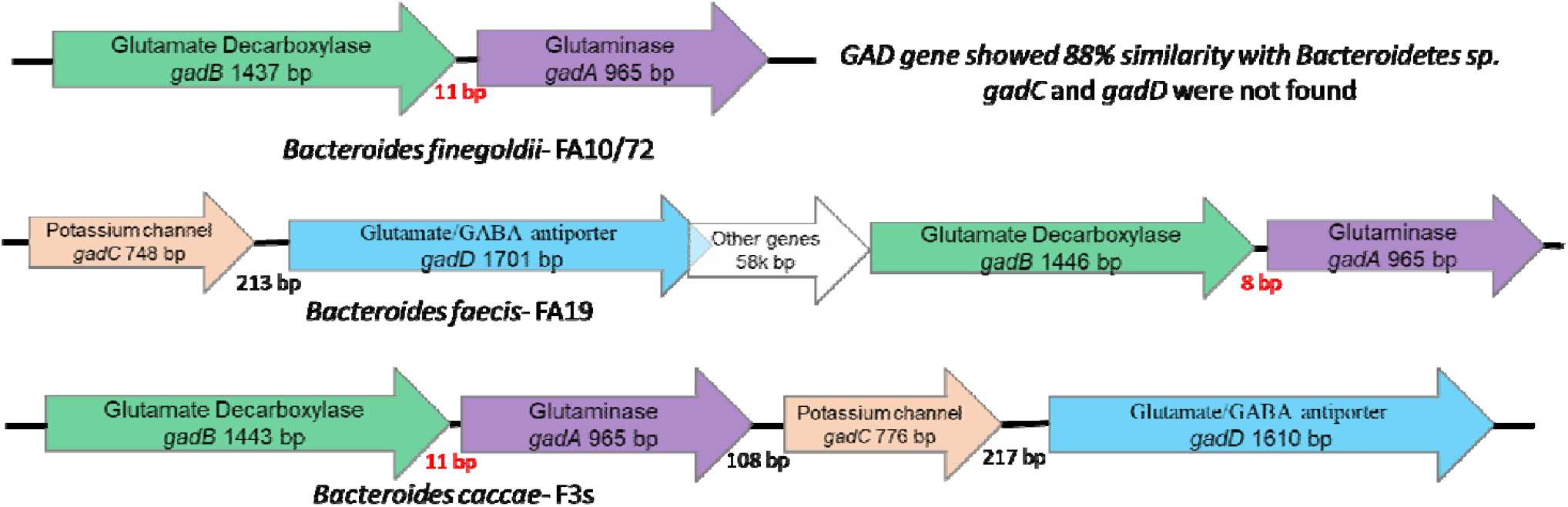
Signature GAD genetic system in high GABA producers

Other important observation amogst these *Bacteroides* strains was the genomic distance between the *gadB*/*gadA* operon and the gadC/K+ channel genes varied considerably from 108 bp to 58k bases difference (**Figure 7**) which could impact gene co-regulation of these two regions (Korbel et al., 2004). Moreover, we observed complete absence of GAD system in *B. Zhangwenhongii* UO.H1054.

Another important characteristic of the *Bacteroides* strains is they harbor set of functionally related genes known as PULs. There are eight genes in PUL, susA-G and susR, including three genes that encode CAZymes which participate the sophisticated breakdown of complex carbohydrates thereby providing health benefitting short chain fatty acids as well a energy. We used a predictive tool to determine number of PULs and their structure amongst these 18 isolates. We observed a great variation in number of PULs and their structural organization amongst these strains (**Table2**).

**Table 2.** Number of PULs identified in each strain.

| Sample Code | Sample ID | Bacterial Isolates | # PUL |
| --- | --- | --- | --- |
| UO.H1001 | BH24/72 | *Bacteroides* sp. | 33 |
| UO.H1004 | BHB5 | *Bacteroides* sp. | 32 |
| UO.H1015 | FA9/72 | *Bacteroides vulgatus* | 51 |
| UO.H1016 | FA12a/72 | *Bacteroides vulgatus* | 48 |
| UO.H1027 | BH19 | *Bacteroides cellulosilyticus* | 94 |
| UO.H1030 | FA9 | *Bacteroides cellulosilyticus* | 90 |
| UO.H1033 | FA14/72 | *Bacteroides dorei* | 58 |
| UO.H1035 | BH27b/72 | *Bacteroides stercoris* | 30 |
| UO.H1039 | BH30/72 | *Bacteroides stercoris* G150 | 31 |
| UO.H1043 | FA3 | *Bacteroides uniformis* |  |
| UO.H1047 | FAB3b | *Parabacteroides johnsonii* | 45 |
| UO.H1049 | FAB6b | *Parabacteroides johnsonii* | 45 |
| UO.H1051 | FA19 | *Bacteroides faecis* | 74 |
| UO.H1052 | FA10/72 | *Bacteroides finegoldii* | 58 |
| UO.H1053 | FA6 | *Bacteroides ovatus* |  |
| UO.H1054 | FA11c | *B. zhangwenhongii* | 74 |
| UO.H2001 | B3s | *B. stercoris* | 38 |
| UO.H2003 | F3s | *B. caccae* | 66 |

### 3.13. Phylogeny of isolates based on 16S rRNA gene and *gadB* genes

Phylogenetic analysis corroborated the uniqueness and highly conserved GAD system genetic arrangement in Bacteroides. Also, we observed congruency in phylogenetic trees based on 16S rRNA and GAD-system gene sequences. Notably, *B. finegoldii* formed a separate lineage as well as *Bacteroides* sp. UO.H1001; UO.H1004 formed a distinct cluster without grouping with other *Bacteroides* strains. This further conforms with the uniqueness of these three strains.

### 3.14. Short Chain Fatty Acid Analysis

Gut bacteria produces 3 major SCFAs: acetate, propionate, and butyrate. Amongst these, acetate production is commonly observed various bacterial classes and is observed in highest concentration in the intestinal lumen whereas propionate and butyrate production are substrate specific (Louis and Flint, 2017). We observed high production of propionic acid (3-5 mM) in most of the strains except *Bacteroides* sp. (UO.H1001, UO.H1004), *B. faecis, B. finegoldii, B. ovatus, B. zhangwenhongii, B. caccae*. Notably, *Bacteroides* sp. (UO.H1001, UO.H1004) showed high concentration of acetic acid (3 mM) in comparison all other strains which showed 1-1.5 mM concentration (**Table 3**). Isovaleric acid was produced by all the strains in 0.8 to 1.9 mM concentrations. Butyric acid was observed in 20-30 uM range whereas Isobutyric acid in 100 to 400 uM range in most of the strains. The production of these SCFAs play a pivotal role in gut integrity as they regulate mucus production, luminal pH, and provide energy to epithelial cells as well as have great impact on immune function.

**Table 3.**
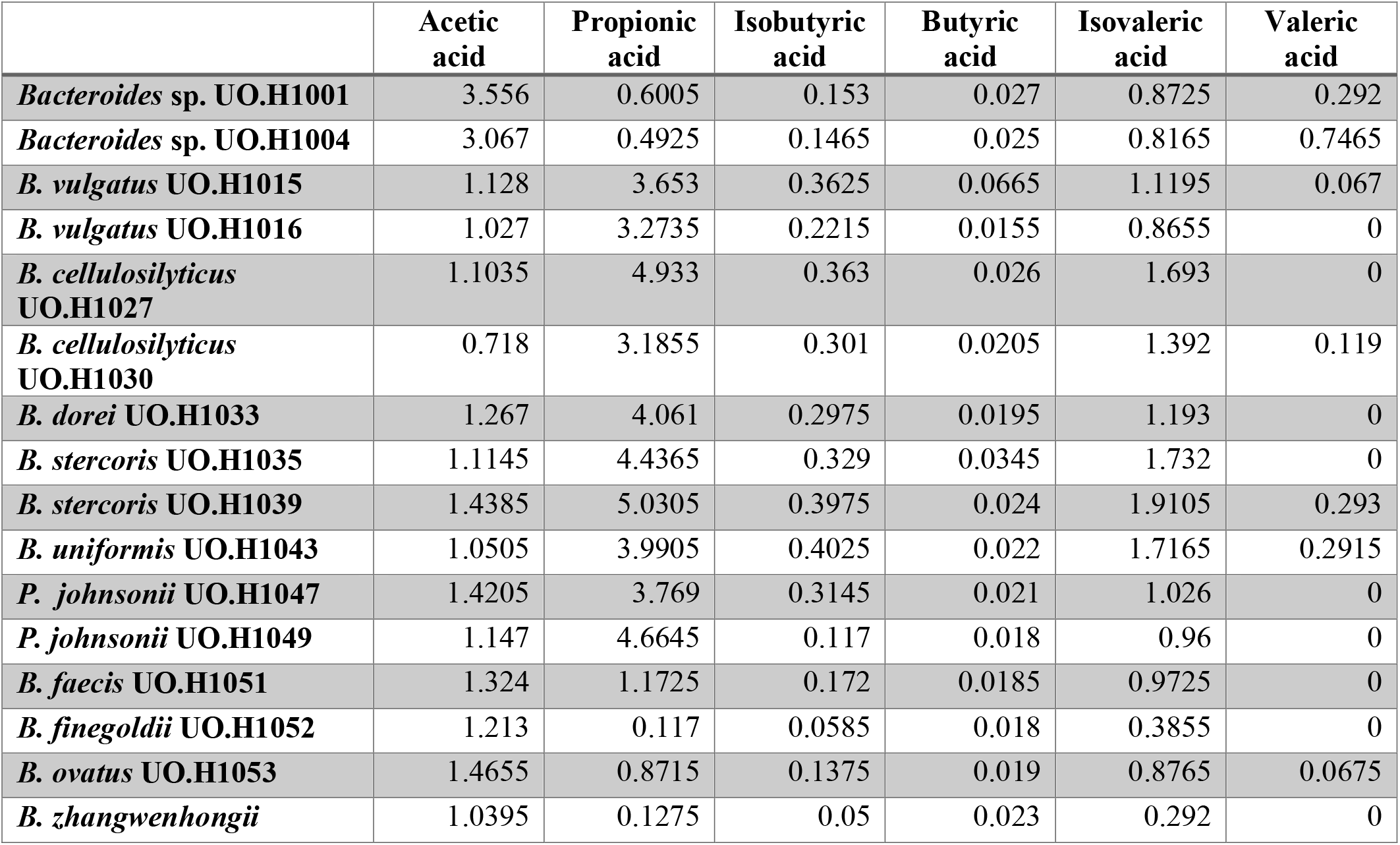
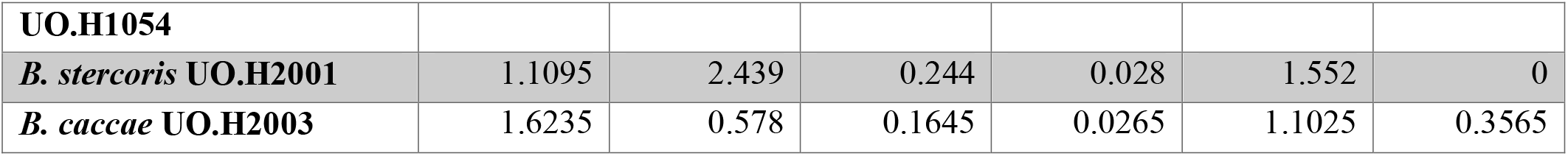
Concentration of Short Chain Fatty Acids (SCFAs) in different Bacteroides strains.

### 3.15. Comparative GABA production analysis

Our *in-silico* analysis showed the presence of diverse *gadA/gadB*/*gadC*/K transporter amongst the 11 *Bacteroide*s species. Thus, we quantified GABA using competitive ELISA approach to see any linkage between GAD genetic system and the GABA production. We observed *B. finegoldii, B. faecis, B. caccae* showed the highest GABA production in 4-7 mM range followed by *B. ovatus* and *Bacteroide*s species in (100-300 μM range) (**Figure 8**). Others showed the presence of GABA in low micromolar concentration or no production. Notably, high GABA producers showed short intergenic region between *gadB* and *gadA* which was not observed in other strains thus indicating possible signature GAD system involved in high GABA production.

**Figure 8.**
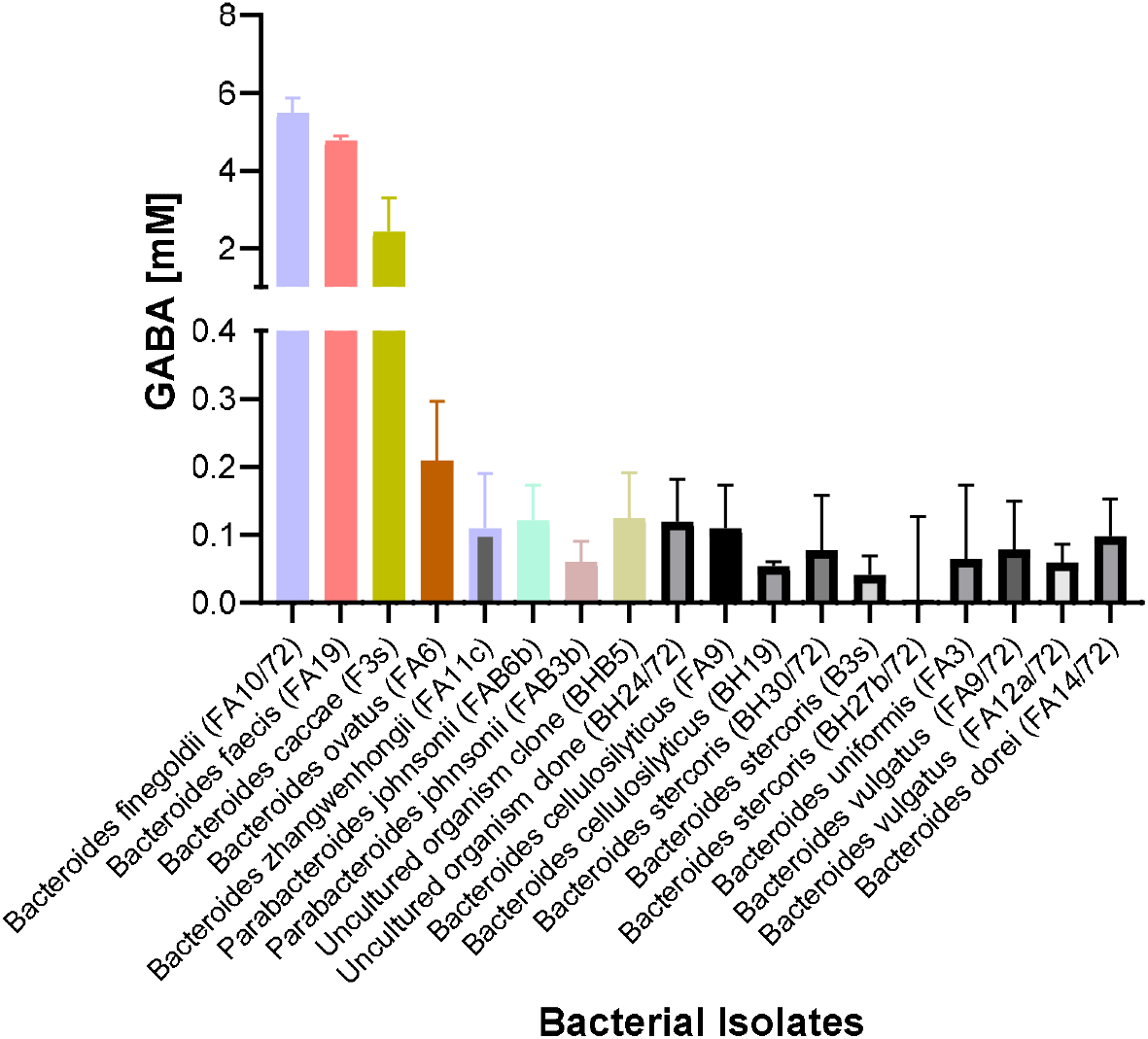
GABA production in 18 representative Bacteroides strains quantified using competitive ELISA.

We extracted EVs from three high GABA producers: *B. finegoldii, B. faecis, B. caccae* and one low GABA producer-*Bacteroides* sp. UO.H1001 and performed the comparative GABA concentration determination amongst these four strains (Supernatant vs EVs). Supernatant showed GABA concentration in 4-7 mM range whereas EVs contained 200-400 μM range (**Figure 9**).

**Figure 9.**
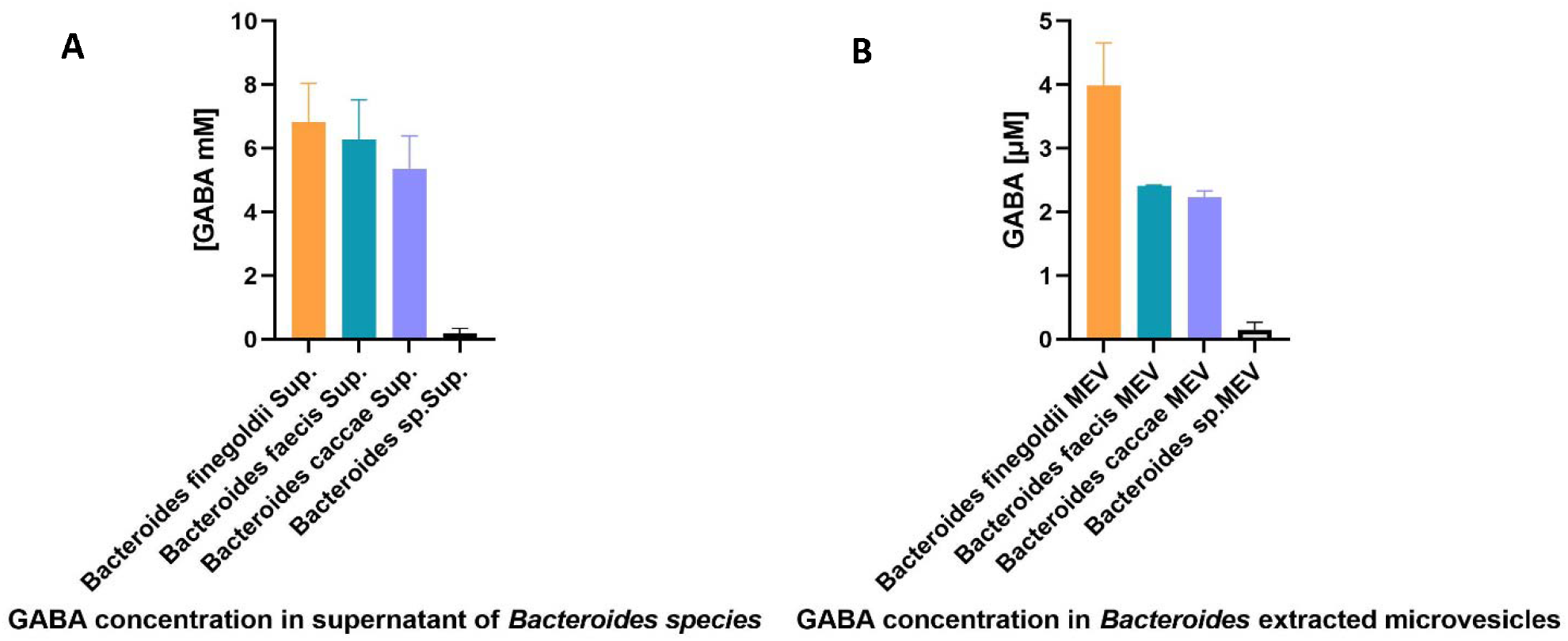
Comparative GABA quantification in bacterial culture Supernatnt (**A**) vs generated EVs (**B**) from high and low GABA producing strains.

### 3.16. Endocytic internalization of MEVs by different cell lines

The endocytic activity of different cell lines that models intestinal transport (CACO-2), secretory (RIN-14B), and blood-brain barrier activities (hCMEC/D3) was evaluated through the addition of Cy7-labeled MEVs to the monolayer culture and compared to the non-treated negative control. CACO-2, RIN-14 B, and hCMEC/D3 cells showed a capacity to internalize labeled MEVs through an endocytic mechanism (**Figure 10**).

**Figure 10.**
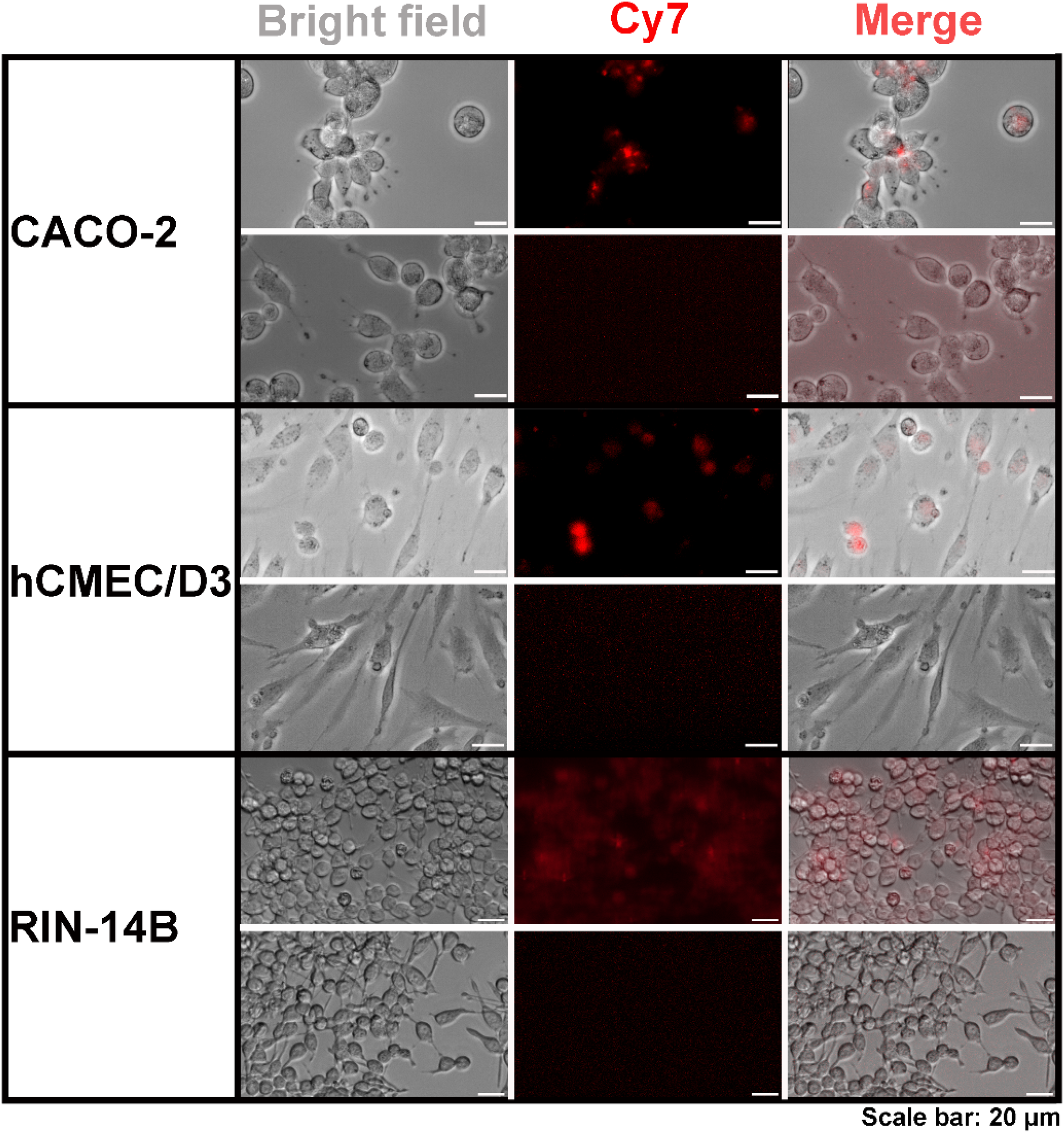
Endocytosis of Cy7-labeled MEVs by various cell types, including CACO-2 (Model of intestinal transport activities), hCMEC/D3 (Model of blood-brain-barrier activities), and RIN-14B (Model of secretory activity).

### 3.17. Transport of MEVs across CACO-2 and hCMEC/D3 cells

FITC-labeled MEVs showed a dose-dependent transport (expressed as paracellular transport (A.U.)) across the intestinal transport model system, CACO-2 cells. The transport of FITC-labeled MEVs (equivalent to 3.6 g stool, 9±1.6 A.U.) was significantly higher after 24 hours as compared to labeled MEVs (equivalent to 1.8 g stool, 4±0.4 A.U., P<0.01) and labeled MEVs (equivalent to 0.9 g stool, 2.8±0.5 A.U., P<0.05) (**Figure 11**). Exposure of CACO-2 cells to different concentrations of FITC-labeled MEVs for 24 hrs significantly decreased the TEER values. (**Figure 11** and **Table 4**). The hCMEC/D3 monolayer cells grown over tissue culture inserts reached a maximum TEER value after 48 hours. Cy7-labeled MEVs showed a dose-dependent transport activity (expressed as paracellular transport (A.U.)) across the blood-brain-barrier model system, hCMEC/D3 cells. The transport of Cy7-labeled MEVs (equivalent to 3.6 g stool, 34.7±0.3 A.U.) was significantly higher after 24 hours as compared to labeled MEVs (equivalent to 1.8 g stool, 17.0±1.2 A.U., P<0.001) and labeled MEVs (equivalent to 0.9 g stool, 7.7±0.3 A.U., P<0.001). In addition, the transport of Cy7-labeled MEVs (equivalent to 1.8 g stool) was significantly higher after 24 hours as compared to labeled MEVs (equivalent to 0.9 g stool, P<0.001). Exposure of hCMEC/D3 cells to different concentrations of Cy7-labeled MEVs for 24 hrs did not alter the TEER values (**Figure 11**).

**Table 4.** TEER values before and after the addition of MEVs to CACO-2 cells as compared to the non-treated negative control.

| FITC-labeled MEVs | 3.6 g stool | 1.8 g stool | 0.9 g stool | Negative control |
| --- | --- | --- | --- | --- |
| TEER values before treatment | 354±5.9 | 270±3.7 | 277±5.8 | 277±4.7 |
| TEER values after treatment | 230±5.5 | 229±4.6 | 249±7.0 | 263±4.9 |
| Statistical significance | $P < 0.001$ | $P < 0.01$ | $P < 0.05$ | $P > 0.05$ |

**Figure 11.**
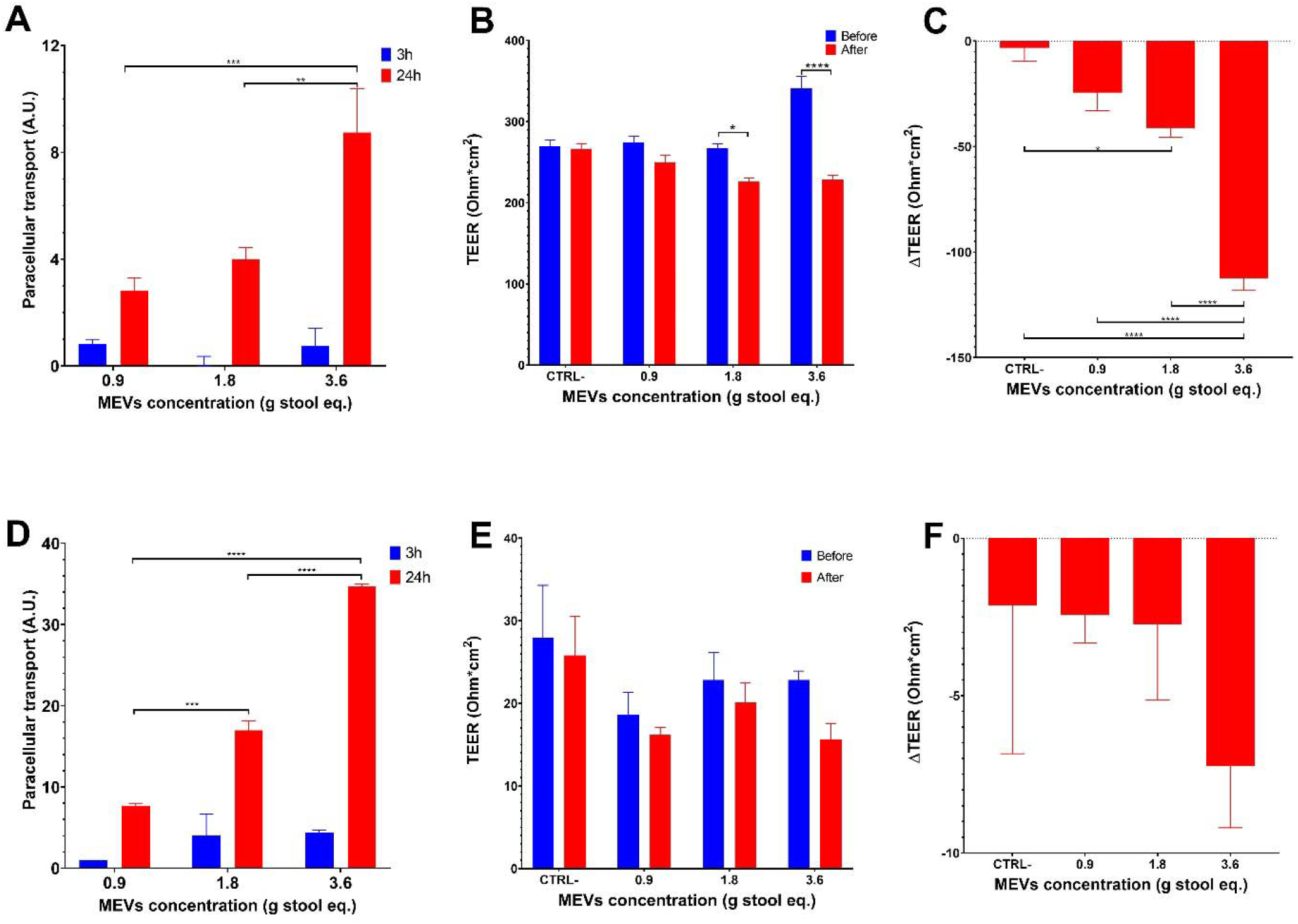
Transport of various concentrations of FITC-labeled MEVs across CACO-2 cells (expressed as paracellular transport (A.U.)). *: P<0.05, **: P<0.01 (A), and TEER values of CACO-2 cells before and after the addition of MEVs (B & C). Transport of Cy7-labeled MEVs across hCMEC/D3 cells (expressed as paracellular transport (A.U.)). ***: P<0.001 (D), TEER values of hCMEC/D3 before and after the addition of MEVs (E & F).

### 3.18. *In vivo* assessment of MEVs biodistribution

The fluorescence intensity expressed as average radiant efficiency was significantly higher in the liver, stomach, and spleen of C57BL6 mice injected intravenously with Cy7-MEVS (equivalent to 12 grams) compared to control mice that received PBS (**Table 5** and **Figure 12**).

**Table 5.** The average radiant efficiency of different organs imaged by IVIS Lumina XR system compared to control.

| Sample | Organ |  |  |
| --- | --- | --- | --- |
|  | Liver* | Stomach* | Spleen** |
| Cy7-MEVs (12 g Stool) | 2.82±0.27 | 1.48±0.17 | 8.86±1.25 |
| Control | 0.19±0.09 | 0.47±0.02 | 0.53±0.18 |
| Statistical significance | P<0.001 | P<0.01 | P<0.01 |

\*Average radiant efficiency $\times 10^7$ , \*\*Average radiant efficiency $\times 10^6$

**Figure 12.**
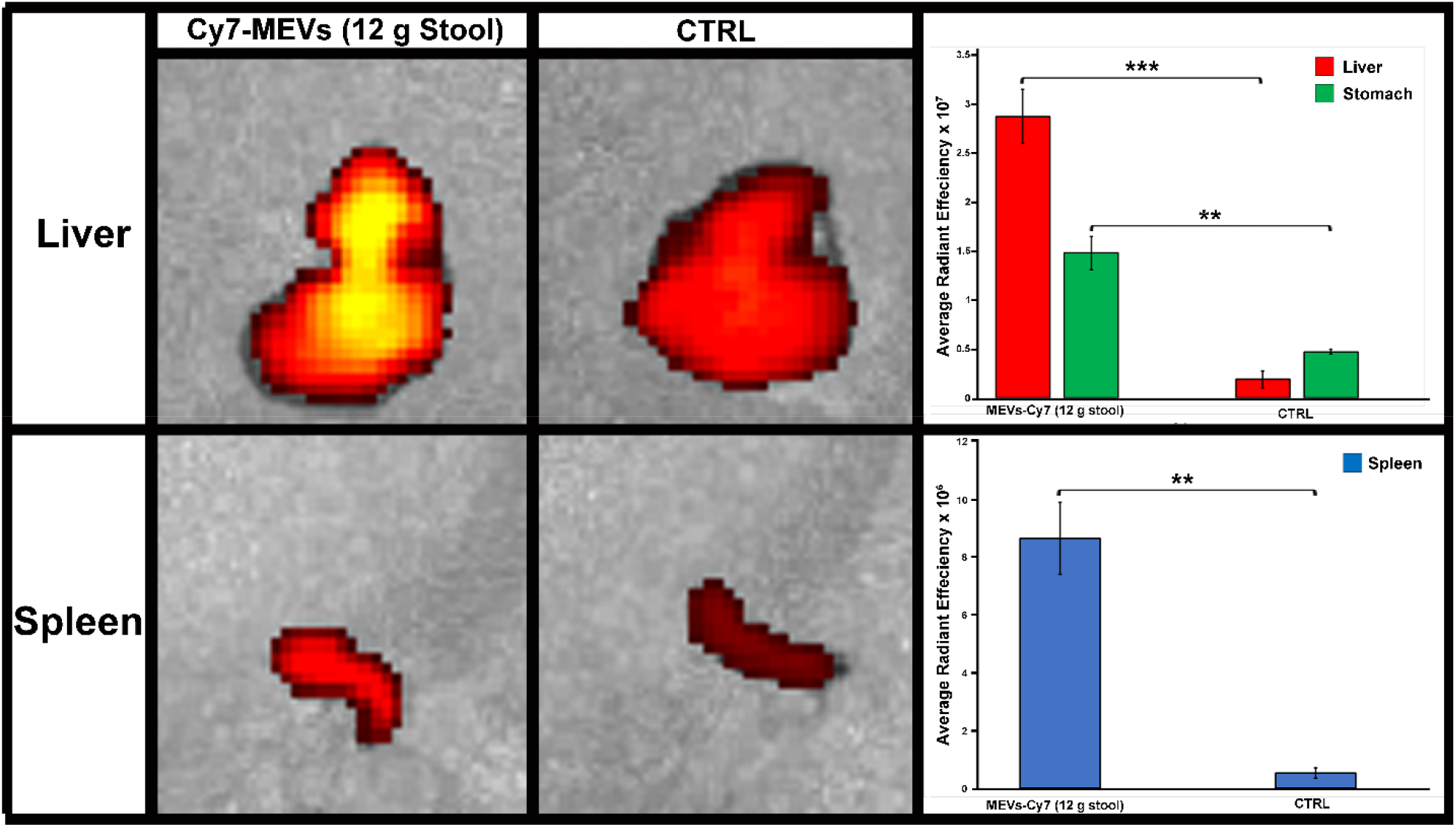
Fluorescent imaging (IVIS Lumina XR) of liver and spleen after intravenous injection of Cy7-MEVs along with average radiant efficiency of different organs after injection of Cy7-MEVS equivalent to 12 g stool as compared to control.

## 4. Discussion

Dysbiosis of intestinal microenvironment has been linked to many health disorders including mental and behavioral disorders. Microbiome based approaches showed promises as potential disease modulating strategies. However, the mechanism by which gut microbiome contributes to these disorders are still hypothetical. In order to achieve a precise modulation of gut microbiome with clinical effectiveness, it is essential to identify the regulatory mechanisms that control the host-microbiome interactions. Recently, microbiome-released extracellular vesicles (MEVs) have emerged as a key delivery mechanism in controlling intestinal microenvironment and bacteria-host communications ^13,48^. MEVs are small membrane-bound phospholipid vesicles that encase a spectrum of biologically active molecules (i.e., proteins, mRNA, miRNA, DNA, carbohydrates, and lipids) that protect them from lytic enzymes and RNases in the extracellular environment^49^ and facilitate their horizontal transfer across both short and distant locations, such as the brain^50,51^. MEVs are involved in numerous processes, such as quorum sensing, biofilm formation, relief of environmental stresses, and host immunomodulation^15^. The production and role of MEVs released by probiotic and commensal microbes in the gut environment are poorly investigated^16,50^, being predominantly examined in pathogenic strains^15,17,18^.

MEVs possess a potential intra- and inter-kingdom signaling mechanism. It is commonly believed that the communication between Gram-negative bacteria and the host is mediated by secreted vesicles, known as outer-membrane vesicles (OMVs)^52^. Gram-positive bacteria have also been reported to generate EVs^53^. In 2013, Kang et al.^54^ reported an important shift in stool MEVs composition compared to the microbiome in a dextran sodium sulfate (DSS)-Induced colitis mice model. While it was unclear whether this dysbiosis was a consequence or a cause of the inflammation, this study illustrates that EVs play a regulatory role in intestinal immunity and homeostasis^54^. Moreover, *Akkermansia muciniphila*-secreted EVs protected mice from developing colitis and lowered the production of the proinflammatory cytokine IL6 in response to an *E. coli* treatment^54^. Additionally, *A. muciniphila-*EVs were reported to induce serotonin secretion in both the colon and hippocampus of mice, suggesting MEVs’potential as signaling molecules in the gut–brain axis^55^. Likewise, *Lacticaseibacillus rhamnosus* GG was shown to produce EVs with immune regulatory activity^56^.

We first reported the capacity of MEVs to cross the epithelial and blood-brain barriers (BBB)^57^. Our results here illustrate the capacity of MEVs to cross and to be endocytosed by intestinal and BBB cell lines. MEVs may cross intestinal barriers and reach distal organs, such as liver and adipose tissues, inducing insulin resistance and glucose intolerance^51^. We found that MEVs have reduced the TEER resistance of CACO-2 cells. This may explain a reported increased level of systemic LPS-positive bacterial EVs in humans with intestinal barrier dysfunction which provides evidence of MEVs’capacity to reach the systemic circulation^58^ and deliver and elicit various immunological and metabolic responses in different organs, including the brain. MEVs were capable to cross the hCMEC/D3 human brain endothelial cell line as a model of human blood brain barrier paracellular permeability^59^. Additionally, MEVs were detected in different organs including liver, stomach, and spleen following IV injection of cy-7-labeled MEVs in C57BL6 mice. In agreement with this, lipophilic EVs, when orally gavaged, injected intravenous or intraperitoneal, were detected in different organs including the brain^60^. This indicates that MEVs that reach the blood circulation are capable to cross the BBB and deliver their cargo to the brain modulating its functionalities. For instance, *Lactiplantibacillus plantarum*-secreted EVs have been reported to suppress stress-induced reduced hippocampal expression of proBDNF and BDNF (Brain-derived neurotrophic factor) in chronic restraint stress (CRST)-treated mice^61^. The same study postulated that *L. plantarum* EVs injected intraperitoneally may reach the brain and induce direct genomic changes on the brain cells^61^.

Our study reveals the presence of a wide range of metabolites embedded in MEVs, including carbohydrates, amino acids, vitamins, and organic acids^57^. Interestingly, We identified many neurotransmitter-related compounds or their precursors inside MEVs, including arachidonyl-dopamine (NADA), gabapentin, and N-acylethanolamines^57^. Dopamine, a representative neurotransmitter in humans, was found in the gut microbiota-derived MEVs as a conjugated form with arachidonic acid. N-acylethanolamines (NAEs) such as palmitoyl-ethanolamide (PEA) and linoleoyl-ethanolamide (LEA) have been reported as effective neuroprotective agent^62,63^. Also, NADA is an endocannabinoid with widespread physiological and pharmacological activities, including modulation of neuropathic pain, inflammatory hyperalgesia, and immune and vascular systems^64^. In accordance with this, a previous report showed that EVs released by *Bacteroides fragilis* include GABA and its intermediates α-ketoglutarate and glutamate as part of their content^65^. The oral administration of *B. fragilis* reduced gut permeability, microbiome dysbiosis, and several behavioral abnormalities in a mice model of autism spectrum disorder (ASD), thus highlighting the potential of microbial interventions in modulating gut microbiome-mediated neurological disorders^66^. Also, *Bacteroides*, a major GABA-producing genus in the gut, was linked with higher levels of serotonin, and myoinositol, which is pivotal in maintaining signaling between the enteric and central nervous system^67^. The relative abundance of *Bacteroides* was negatively correlated with depression-associated brain signatures^68^, indicating a significant role of microbiome-secreted GABA in brain functionality. Likewise, Mason et al.^69^ have reported depletion of *Bacteroides* in depression and anxiety. Finally, several lipid species in our dataset are conjugated with high unsaturated fatty acids (HUFAs) such as linoleic acid and arachidonic acid. HUFAs are key molecules for the development, maintenance, and performance of the nervous system as well as brain functionality through improving the fluidity of the cell membrane^70^. Collectively, this indicates that gut microbiota-derived MEVs contain neurotransmitter-like molecules proposing MEVs as a signalling shuttle in the gut microbiota-brain axis.

Neurochemicals, such as GABA, serotonin, dopamine, or their precursors and derivatives, are microbially metabolized by gut commensals and being considered major modulators of the gut environment, including the enteric nervous system^71^. We found 3,4-dihydroxymandelic acid (DOMA), a metabolite of norepinephrine as a part of MEVs’content. Sule and coauthors have reported that 3,4-dihydroxymandelic acid is produced by the metabolic activity of *E. coli*^72^. Also, *Bifidobacterium dentium*, a GABA-producing bacterium, modulated sensory neuron activity in a rat fecal retention model of visceral hypersensitivity. Besides, GABA was detected in the cytoplasm and brush border of epithelial cells in the rat jejunum and colon^73^. The exposure of GABA to epithelial cells selectively stimulated MUC1 expression in isolated pig jejunum^74^, and increased the expression of tight junctions and transforming growth factor beta (TGF-β)^75^ while decreased IL-1β-mediated inflammation *in vitro*^75^, providing a protective effect against the disruption of the intestinal barrier. Importantly, GABA has also been identified as an essential growth factor that solely can induce the growth of unculturable gut microbes^68^. Together, this indicates that the metabolites imbedded in MEVs may also modulate the gut microenvironment.

Overall, our results provided pioneering and significant insights into MEVs’capacity to transfer neuroactive metabolites to the host intestine and other organs, including the brain, filling some of the gaps in knowledge of the mechanisms underlying microbiome-gut-brain interactions.

## Acknowledgments

This study was supported by a grant from the Weston Family Foundation, through its Weston Family Microbiome Initiative and an NSERC-Discovery grant RGPIN-2018-06059. S.S., N.E.B., and J.Y. were supported respectively by the Nutrition and Mental Health Master’s and doctoral Scholarship and a postdoctoral fellowship from the University of Ottawa.

## Ethical Statement

The study was conducted following the Declaration of Helsinki and approved by the University of Ottawa Research Ethics Board and Integrity (protocol code H-02-18-347 and approved on 29 July 2019).

